# CADD-SV – a framework to score the effects of structural variants in health and disease

**DOI:** 10.1101/2021.07.10.451798

**Authors:** Philip Kleinert, Martin Kircher

## Abstract

While technological advances improved the identification of structural variants (SVs) in the human genome, their interpretation remains challenging. Several methods utilize individual mechanistic principles like the deletion of coding sequence or 3D genome architecture disruptions. However, a comprehensive tool using the broad spectrum of available annotations is missing. Here, we describe CADD-SV, a method to retrieve and integrate a wide set of annotations to predict the effects of SVs.

Previously, supervised learning approaches were limited due to a small number and biased set of annotated pathogenic or benign SVs. We overcome this problem by using a surrogate training-objective, the Combined Annotation Dependent Depletion (CADD) of functional variants. We use human and chimpanzee derived SVs as proxy-neutral and contrast them with matched simulated variants as proxy-pathogenic, an approach that has proven powerful for SNVs.

Our tool computes summary statistics over diverse variant annotations and uses random forest models to prioritize deleterious structural variants. The resulting CADD-SV scores correlate with known pathogenic and rare population variants. We further show that we can prioritize somatic cancer variants as well as non-coding variants known to affect gene expression. We provide a website and offline-scoring tool for easy application of CADD-SV (https://cadd-sv.bihealth.org/).

## Introduction

In the light of recent advances in the field of structural variant (SV) detection and the study of regulatory domain architectures, phenotypic effects of SVs in humans moved into the focus of research [1–4]. SVs can be deletions, duplications, insertions, translocations or inversions and often span multiple kilobases of sequence in the genome. Due to their size, they have the potential to cause significant phenotypical effects and are therefore relevant for clinical genetics [1,3,5,6]. While SVs affecting the expression of whole genes or exons are still the research focus, effects of non-coding DNA sequence alterations are of high interest. These variants are especially hard to predict, as our understanding of such regions lags behind coding annotations [7]. In comparison to pathogenic variants (e.g. frame shift mutations or disruption of transcription factor binding) caused by single nucleotide variants (SNVs), structural variants have a higher potential to affect the regulatory architecture of the genome. Thus, the functional characterization of SVs may help us to understand unexplained disease phenotypes and contribute to our understanding of regulatory mechanisms.

Recent advances in the study of regulatory genome architectures provided evidence along these lines and already shed light on previously unexplained human disease conditions [8,9]. The most relevant examples are improved Hi-C protocols to study genome architecture [10], the experimental annotation of enhancers and enhancer-promoter links [11], mapping of multiple epigenetic features across many cell-types [12], but also methods to test the regulatory potential of sequences in high-throughput [13–16]. All these advances provide a basic understanding of topological domain structures, regulatory elements and other fundamental mechanistic insights like enhancer hijacking [17,18]. However, wider understanding of how SVs link to phenotypic alterations and therefore human diseases remains poor.

SV identification and annotation lags behind SNV and InDel annotation as SVs often exceed the size of common read-lengths, are difficult to align, fall within repetitive regions or can be of complex structure [19]. In addition, various factors may contribute to pathogenicity or molecular effect in these regions as structural rearrangements can affect primary gene structure, chromatin architecture, DNA accessibility and tissue-specificity of regulatory elements and genes. Further, the putatively different mechanisms of phenotypic effects of deletions compared to insertions or duplications complicates a generalized approach for variant effect prediction as the effect can be mediated by copy number alterations of redundant or unique genomic sequence, positional effects or rendering functional DNA dysfunctional. Capturing all possible disease relevant mechanisms mediated by structural variants remains challenging.

While various tools are available for ranking SNVs and small insertion/deletions (InDels), very few tools can score structural variants. Therefore, it remains very difficult to assess SV effects on phenotype and disease, with many different ad-hoc approaches being applied. Existing tools like SVScore [20] or TAD-Fusion [21] focus on individual features such as the presence of deleterious SNVs (mostly in coding regions) which are overlapping the SV or focus specifically on boundary element reshuffling by a novel SV, respectively. AnnotSV [22] annotates the structural variant and categorizes pathogenicity depending on overlap with known pathogenic SVs. SVFX [23] provides a framework for training specific models, but does not allow the direct application to novel variants. At this stage, no tool combines ease of use with a comprehensive set of annotations, including the prioritization of disease effects from genome architecture alterations.

Further, SV data sets of sufficient size and curation that can be used to apply Machine Learning approaches for the identification of relevant annotations or for their integration are not easy to obtain. Clinically relevant SV sets [24], i.e. pathogenic and benign variants, are small in number, biased towards very large SVs and tend to overlap well studied disease genes. Here, we present a novel Machine Learning approach (CADD-SV) to score the effects of SVs by choosing an unbiased and sufficiently large training dataset derived from species differences. For training and application of our models, we implement fast SV annotation and data integration of diverse genomic features (incl. regulatory and 3D architecture). We validate model performance on independent datasets of germline and somatic SVs. Our tool enables an easy application to novel SVs and its normalized features values give insights into potential underlying phenotypic effects.

## Materials & Methods

### Training dataset

We use evolutionarily fixed chimpanzee and human derived SVs from Kronenberg et al. [25]. We refer to the human and chimpanzee deletions and insertions from this set as proxy-neutral or proxy-benign. A set of randomly distributed SVs over the human autosomes was obtained by shuffling the ape SVs matched in length and number (within coordinates considered alignable by Kronenberg et al.). We refer to this set as proxy-pathogenic. To compare these SVs with those in ClinVar [24], we annotated them with the distance to the next start codon, pLI and haploinsufficiency scores (Supplemental Figure 1). We use sets of variants derived from human and chimpanzee to score different SV types. For novel human deletions, we chose the chimp deletions to model the span and human deletions to model the SV flank. Respective annotations are present along the range of chimpanzee deletions in the human genome build, while they are absent for derived human deletions. Similarly, to score insertions, we use the derived human insertions to model the flank and the chimpanzee insertions to model the site of an insertion. Duplication sites are modeled by the chimpanzee deletion model for span and human insertion model for the flank, as the span of duplications contains known sequence most similar to the one found in annotated deletion sequences.

### Feature annotation and transformation

We obtained a set of 127 continuous human derived features (see Supplemental Table 1) ranging from species conservation, distance to gene model hallmarks, over to genome architecture features such as the directionality index derived from Hi-C datasets. We use customized bash and R scripts to annotate the contrasting SV sets using bedtools [26]. All features are Z-score (mean 0, variance 1) transformed using 20,000 randomly selected SVs of the same-type from healthy individuals reported in the gnomAD-SV release v2.0 [2]. All SVs are annotated over the span of the primarily affected sequence (span) as well as 100 bp up- and downstream of the site of the structural rearrangement (flank). From the different annotations, we create summary statistics and transformations as model features. These are summarized in Supplemental Table 1. The annotation framework automatically retrieves the features from primary annotation sets using the workflow management system Snakemake [27]. It tabulates results in a BED-like format that is used in the CADD-SV model. Missing values are imputed with zeros.

### Models

We trained logistic regression and random forest classification models contrasting proxy-benign and proxy-pathogenic training datasets. Models are trained in R (v3.5.1) for the SV spanning sequence for deletions and duplications and the site of integration for insertions (span), as well as 100bp up- and downstream of the reported breakpoints (flank, see Supplemental Figure 2). For logistic regression, we use the R generalized linear model implementation and for random forests the package “randomForest” [28]. For random forests, we limit the number and depth of the decision trees based on a hyperparameter search (Supplemental Figure 3; explored ranges for ntree = {25, 50, 75, 100, 200, 500, 1000}, nodesize = {10, 50, 100, 250, 500, 1000}, maxnodes = {10, 50, 100, 250, 500, 1000}, while one parameter was optimized, the other parameters were set to 100). We randomly withheld 10% of the annotated SVs as hold-out and assessed model performance metrics using the R Package PRROC [29].

### CADD-SV scoring and normalization

Each novel SV is scored using the span and flank models of the respective SV type. To make models comparable, both span and flank scores are ranked according to the score distribution of the same type of SV identified in gnomAD-SV release v2.0 [2]. To capture pathogenicity mediated by the affected sequence as well as sequence context, the higher ranking (more pathogenic) score is reported as the final score of the event. As a result, the CADD-SV score equals the rank percentile of the novel SV in the distribution of 20,000 SVs of the same type identified in healthy individuals. Values range from zero to one, where a value of one signifies that no SV with a higher score was detected in the healthy cohort.

### Model validation

Pathogenic and benign annotations for clinical SVs [24] were downloaded from ClinVar (https://www.ncbi.nlm.nih.gov/clinvar) on June 24th, 2021. Only variants with pathogenic or benign labels of at least 50 bp and annotated as deletion (pathogenic n = 3262, benign = 33), duplication (pathogenic n=82, benign n = 4) or insertion (pathogenic n = 78, benign n = 18) are considered. Further, to increase the number of pathogenic insertions, unique pathogenic insertions (n = 39) reported in Hancks et al.[30] and Gardner et al [31] were added. Area under the receiver-operator curve (AUROC) metrics are calculated using the PRROC R-package [29].

Germline SVs identified from healthy individuals over various populations [2] were downloaded from gnomAD-SV release v2.0 (https://gnomad.broadinstitute.org/downloads). Allele frequency values as well as common and ultra-rare SVs are determined across all available populations. Common variants are defined as minor allele frequency greater 0.05, ultra-rare variants are defined as singletons. To show the clinical benefit of prioritization of SVs using CADD-SV, we use 1000 Genome genotyped SVs [32] and add one (randomly selected) labelled pathogenic SV as found in ClinVar into the reported set of individual specific SVs. From the 1000 Genome events annotations, we consider Alu and Line1 SVs to be insertions. We report the rank of the pathogenic SVs within the occurring SV sets.

Somatic SVs (n = 95,749) from cancer patients were obtained from the International Cancer Genome Consortium [33] at https://dcc.icgc.org/api/v1/download?fn=/PCAWG/consensus_sv/final_consensus_sv_bedpe_passonly.icgc.public.tgz. In addition, insertions reported in cancer genomes were taken from Qian et al. (n = 18) [34]. To assess the performance of CADD-SV beyond coding regions, we use non-coding SVs (n = 687) that are known to impact human gene expression in data from the GTEx consortium [1].

### SV scoring tools

CADD-SV performance on various validation sets was compared to existing tools SVScore [20], AnnotSV [22] and, for deletions, the TAD-fusion-score [21] using standard parameters. As SVScore and TAD-Fusion score were not designed for the current genome build GRCh38, UCSC liftover [35] was used to transfer SV coordinates and respective scores.

### Implementation

Novel SVs can be scored with a pipeline implemented in Snakemake [27], using conda [36] for dependency management. The source code for the framework is available for download on GitHub (https://github.com/kircherlab/CADD-SV/). A webservice (https://cadd-sv.bihealth.org/) allows for online scoring of SVs in BED format as well as for obtaining results for different human genome builds (GRCh38; NCB16 & GRCh37 through automated coordinate liftover). In addition, pre-scored variants from cohorts such as gnomAD or ClinVar with all annotations can be queried online including all features. For better interpretability, feature outlier values are color-coded based on their Z-scores.

## Results

### Large and unbiased training data set

Machine Learning methods strongly rely on the quality of training datasets to yield meaningful predictions. Using clinical databases such as CinVar or HGMD to curate an annotated training dataset is challenging for SNVs or small InDels, where it requires a careful matching of pathogenic and benign variants in genomic regions and effect classes [37,38]. This seems currently impossible for SVs. The ClinVar dataset [24] is very sparse for SVs, i.e. only few (3,262 deletions, 82 duplications and 78 insertions) and mostly very large SVs (mean size of 106 kb for deletions) are being annotated. This is insufficient for an insightful training dataset, especially as population SVs are much smaller (mean size of 7.4 kb). Further, when compared to large population SV sets [39], strong biases towards high effect variants clustering around well studied genes are apparent (Supplemental Figure 1). Therefore, we opt for an unbiased evolutionary set of SVs obtained from comparisons in the great ape lineage [25]. A key strength of this approach is that the model is trained on a larger training set of 19,113 deletions and 26,823 insertions and duplications that does not suffer from the ascertainment bias inherent to curated sets (Suppl. Figure 1).

This is motivated by the Combined Annotation Dependent Depletion (CADD) framework, an approach that has proven powerful in the interpretation of SNVs and short InDels [40]. In CADD-SV, we assume that millions of years of purifying selection removed SVs that are deleterious, i.e. have a negative impact on human or chimpanzee reproductive success. Thus, fixed SVs in humans or chimpanzees can be classified as proxy-neutral. In contrast, variants of the same size randomly drawn from the human genome are likely to contain a significant number of deleterious variants by chance. While many of the random variants will be neutral, an unknown but considerable fraction would likely be deleterious. For simplicity, we refer to these variants as proxy-deleterious. The contrast between the proxy-neutral and proxy-deleterious variant sets, i.e. the relative paucity of deleterious, phenotypically influential genome alterations in the proxy-neutral set and the resulting differences in their annotation features, is the core characteristic of what we then model as SV deleteriousness (Fig. 1).

**Figure 1:**
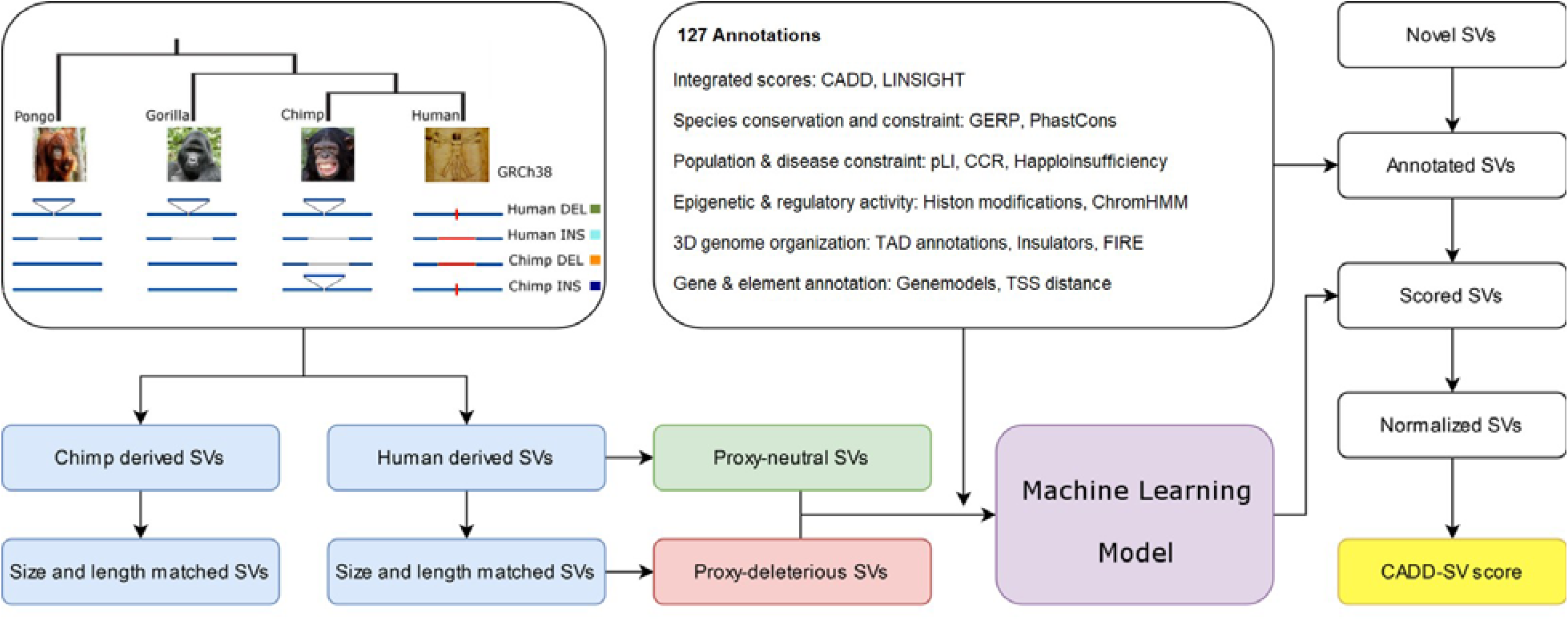
Depiction of the CADD-SV model and workflow. Human and chimpanzee derived SVs from Kronenberg et al. [25] are used as proxy-neutral training dataset. Size and length matched simulated variants are used as proxy deleterious training dataset. Next, various informative features are annotated and transformed (see Methods and Suppl. Table 1) across span or flank of the variants to train Random Forest classifiers. Models are used to score user provided novel SVs. For this purpose, variants are annotated, features transformed and models applied. Raw model scores are ranked among 20,000 gnomAD SVs of the same type and the relative rank returned as the output CADD-SV score.

### Annotating Structural Variants

We wanted to integrate diverse annotations into predictive, genome‐wide models for identifying variants of phenotypic effect. While many annotations are readily available for SNVs, informative and computational efficient statistics need to be created to summarize annotations over the span of SVs. Further, distance measures can retain information about the vicinity of the impacted DNA sequence. For this purpose, we developed an automated SV annotation pipeline using the workflow management system Snakemake [27] that combines bedtools [26] with customized bash and R scripts. We integrate not only coding information such as gene models but also a wide variety of regulatory annotation retrieved from ENCODE [12] such as histone modifications or DNA accessibility. In addition, we make use of functional and evolutionary scores [37,38,41,42] as well as information about the 3D architecture of the genomic region derived from Hi-C experiments [43–45].

All SVs are annotated over the full span of the event as well as 100 bp up- and downstream (Suppl Figure 2). For insertions, the span of novel SVs only contains the site of integration and CADD-SV does not derive features from the inserted sequence. While deletions directly remove putatively functional sequence, insertions and duplications interfere with functionality by integration of additional sequence, e.g. disrupting regulatory interactions by increasing distance or introducing frameshifts into coding sequence. We incorporate this in the CADD-SV modelling by deriving features from the deleted sequence (span), annotating the context of the SV (flank) and including distance features in the model. Across SV ranges, we mostly annotate max values, mean values and the amount of high impact values above the top 90^th^ percentile. Additionally, span and flank models use genomic distances to certain feature coordinates (e.g. genes, exons, and enhancers). All features and their transformation are described in Supplemental Table 1. To ease later interpretation of feature impact, all features are Z-score transformed using the annotation distribution of the same type of SV from healthy individuals reported in gnomAD [39].

### Modeling and hold-out set performance

SV mediated pathogenicity depends on the type of SV. We implement separate models for deleted, inserted, or duplicated sequence. Due to the lack of training data for inversions and translocations, we can currently not train models for these variant types. Using the described training data sets, we train four types of models (Fig. 1). We train models of human-derived deletion (human DEL) and insertion (human INS) events against respective sets of equally sized events drawn across the genome. Further, models based on chimp insertion (chimp INS) and deletions (chimp DEL) events are trained. Here, we project the events onto the human reference sequence and use the human annotations. While the human events are also manifested in the human reference, the chimp events allow us to use human annotation unimpaired by an actual SV event. Hence, chimp DEL models are similar to how we would score new events observed in an individuals' genome aligned to the human reference sequence. In contrast, no annotation for human derived deletions can be obtained over the span of the deletion as experimental readouts and conservation score are not available for the missing sequence. Similarly, chimp INS provide an insertion model based on events that did not impair human annotations or biochemical readouts.

To score novel SVs in the human genome we exploit this relationship by training the span of novel deletions with the chimp DEL set and train the sequence 100bp up- and downstream of the breakpoints using the human DEL set. As the inverse applies for insertions and duplications, i.e. chimpanzee insertions do not span sequence in the human genome build while human derived insertions do, we use the chimp INS set for the insertion site and the human INS set for the up- and downstream sequence. Duplications are scored using the full sequence span of the duplicated locus, hence using the chimp DEL model for the span and human INS model for the up- and downstream sequence. The final score is calculated from the maximum (more deleterious) value of both models applied after being ranked compared to the score distribution of the same type of SV reported in healthy individuals.

We trained both logistic regression models as well as random forest models. We note that the latter show increased hold-out performance as well as validation set performance (Suppl. Figure. 3) and we only describe the random forest models here. The hold-out shows that all four model types differentiate between the proxy-benign and proxy-pathogenic sets (Fig 2A). Considering the anticipated mislabeling in our training data, specifically in the randomly drawn SVs as described above, the hold-out performance will however not be representative for our models' performance in scoring actual pathogenic versus benign variants. Here, we only look for a non-random model performance and the relative ranking of the models. For better interpretation, we also normalize the final score distribution in a healthy population cohort (gnomAD-SV) separately for each type of SV. The score distribution for the hold-out data is available in Fig. 2B for the proxy-pathogenic and proxy-benign SV sets. We see a significant shift with a bimodal distribution in the proxy-pathogenic variants, with the smaller mode corresponding to the potentially pathogenic variants in the randomly drawn set.

**Figure 2:**
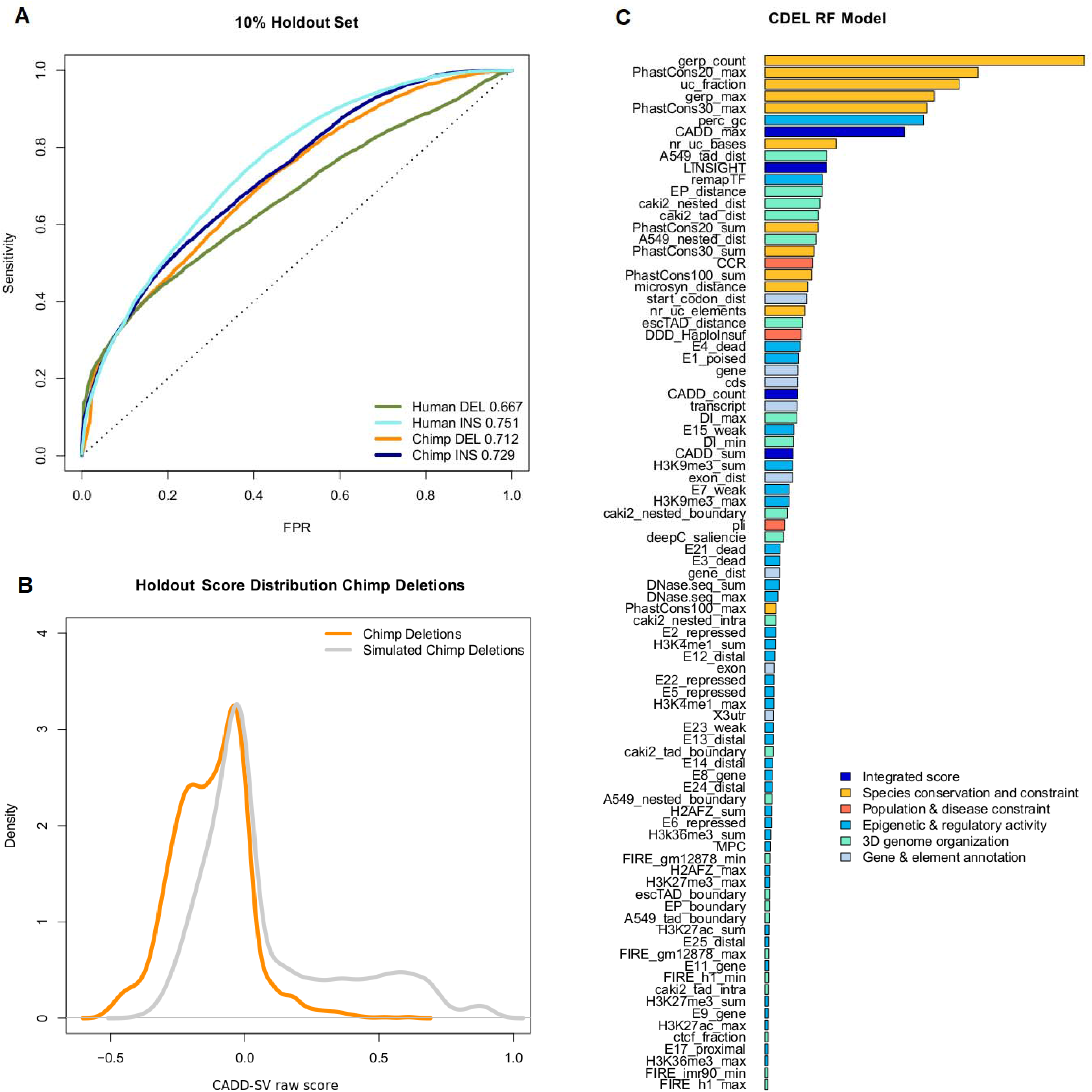
Performance of Random Forest models trained on proxy-deleterious and proxy-benign SVs. **A)** All models show a non-random separation of the two classes in a random 10% holdout. Performance is measured as sensitivity over false positive rate (FPR). Note that all training datasets contain a high amount of mislabeled SVs, as a majority of proxy-deleterious SVs is likely to be neutral. **B)** Model predictions of the Chimp DEL model are shifted towards high impact SVs in the simulated set of chimpanzee deletions. **C)** Representation of feature importance in the chimpanzee deletion (chimp DEL) Random Forest model. Note that proxy-pathogenic and proxy-benign sets are length matched and that length is not used as an explicit feature. Most important contributions come from species conservation (e.g. GERP, PhastCons) but also from integrated scores (i.e. CADD or LINSIGHT). Epigenetic features as well as 3D genome architecture features such as the Directionality Index derived from Hi-C data also contribute to the most informative features of the models. For a full list of features and explanation of their naming, see Suppl. Table 1.

### Feature contributions

We analyzed feature contributions in our random forest models using the R package randomForest [28]. To ease interpretation, we categorized model features into six groups (“Integrated scores”, “Species conservation and constraint”, “Population and disease constraint”, “Epigenetic and regulatory activity”, “3D genome organization”, “Gene and element enrichment”; Supplemental Table 1). Models benefit highly from features in the groups of “Species conservation and constraint” (incl. GERP, PhastCons, PhyloP scores) and "Integrated scores" (i.e. summaries of CADD SNV and LINSIGHT scores) in differentiating between the contrasted SV sets. Regulatory annotations as well as 3D genome architecture features contribute to a smaller extent but are present within the top 20 most important features of all models (e.g. ReMap transcription factor occupancy, TAD annotations, enhancer-promotor links and chromHMM states). Distance features (such as distance to coding sequence) are particularly prevalent in the human DEL flank model, where for a reference altered by the deletion event these features become informative. Major feature contributions of the chimp DEL model are presented in Figure 2C, for all models feature importance is available in Suppl. Figures 5-8.

### Independent Validation Datasets

To validate the general applicability of the framework, we use multiple lines of evidence (Fig. 3A) to substantiate the results of the hold-out performance. We look at known pathogenic variants from ClinVar (Fig. 3B, 3D-F), we show that SVs occurring in healthy populations are under negative selection and therefore high CADD-SV scores enriched for singletons events (Fig. 3C), we analyze variants from the International Cancer Genome Consortium (Fig. 3D-F), and SVs affecting gene expression (Fig. 3D-F). Thereby, we show that CADD-SV can be used to prioritize both pathogenic germline and somatic structural variants.

**Figure 3:**
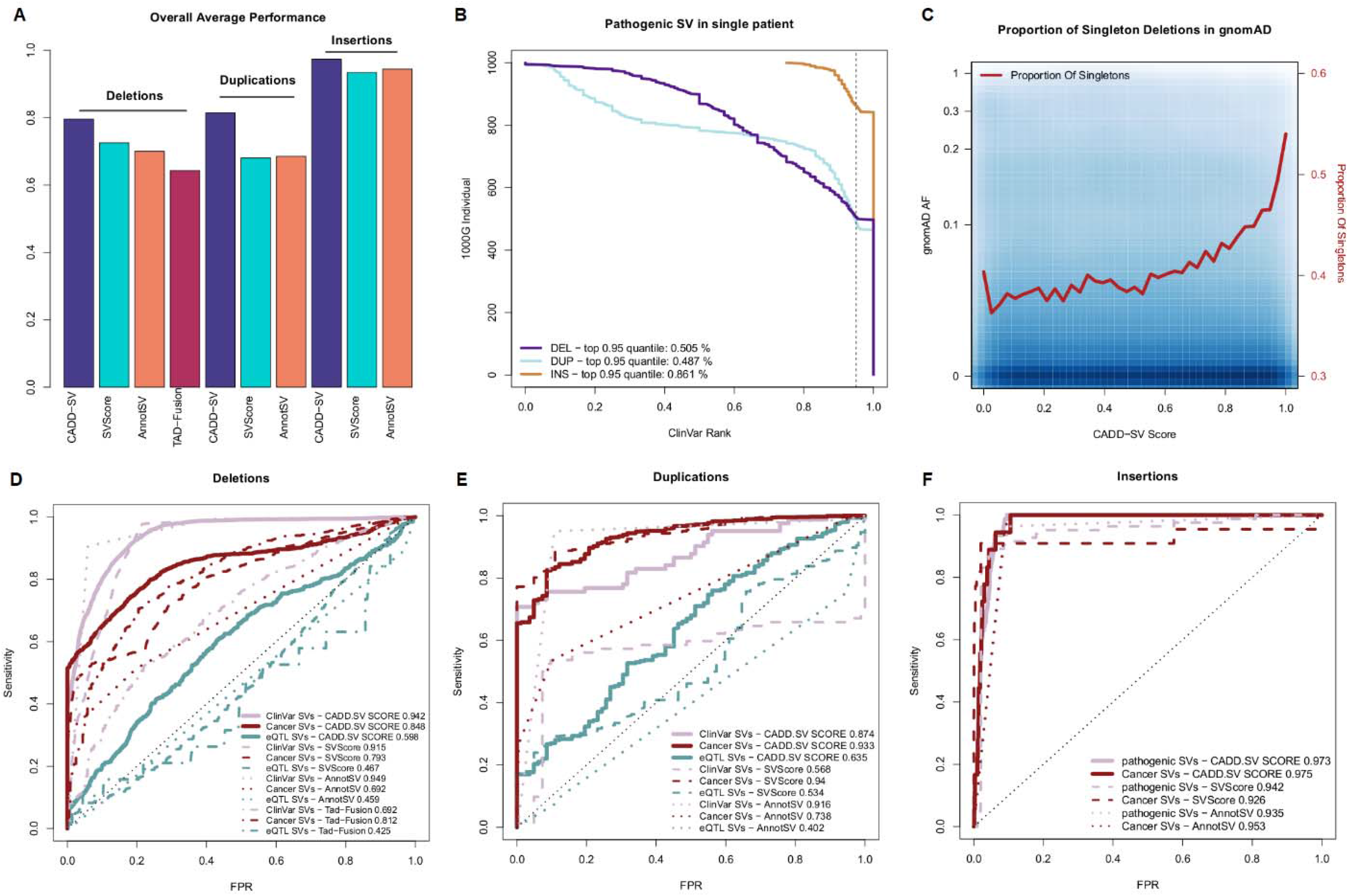
Validation set performance of the Random Forest models. **A)** Summary of the performance of CADD-SV scores compared to SVScore, AnnotSV and TAD-Fusion scores across three validation sets (pathogenic variants, cancer variants and putative eQTL SVs) for deletions, duplications and insertions. **B)** Rank of ClinVar pathogenic SVs added to SVs of healthy individuals from the 1000G projects. CADD-SV prioritizes the pathogenic SVs over the SVs in a single patient, scoring pathogenic variants in the top fifth percentile of deletions, duplications and insertions at 50.5%, 48.7% and 86.1% respectively. **C)** CADD-SV score distribution as a function of gnomAD allele frequency. Higher CADD-SV values represent an increased likelihood to be deleterious. In the deleterious tail of the score distribution, there is an excess of singletons (shown in red; bin-size 0.025), which might hint towards negative selection against deleterious deletions. **D-F)** CADD-SV performance of various validation sets compared to common gnomAD SVs (AF >= 0.05). Performance is measured as sensitivity over false positive rate (FPR). CADD-SV is able to identify ClinVar pathogenic deletions (pale red) as well as deletions reported in the ICGC cancer cohort (dark red) from SVs in gnomAD. Further, CADD-SV can identify non-coding SVs that are associated with differences in gene expression (turquoise). CADD-SV scores (solid lines) are compared to SVScore (dashed lines), AnnotSV (dotted lines) and TAD-Fusion (dashed and dotted lines) for deletions **(D)**, duplications **(E)** and insertions **(F)**.

**Figure 3:**
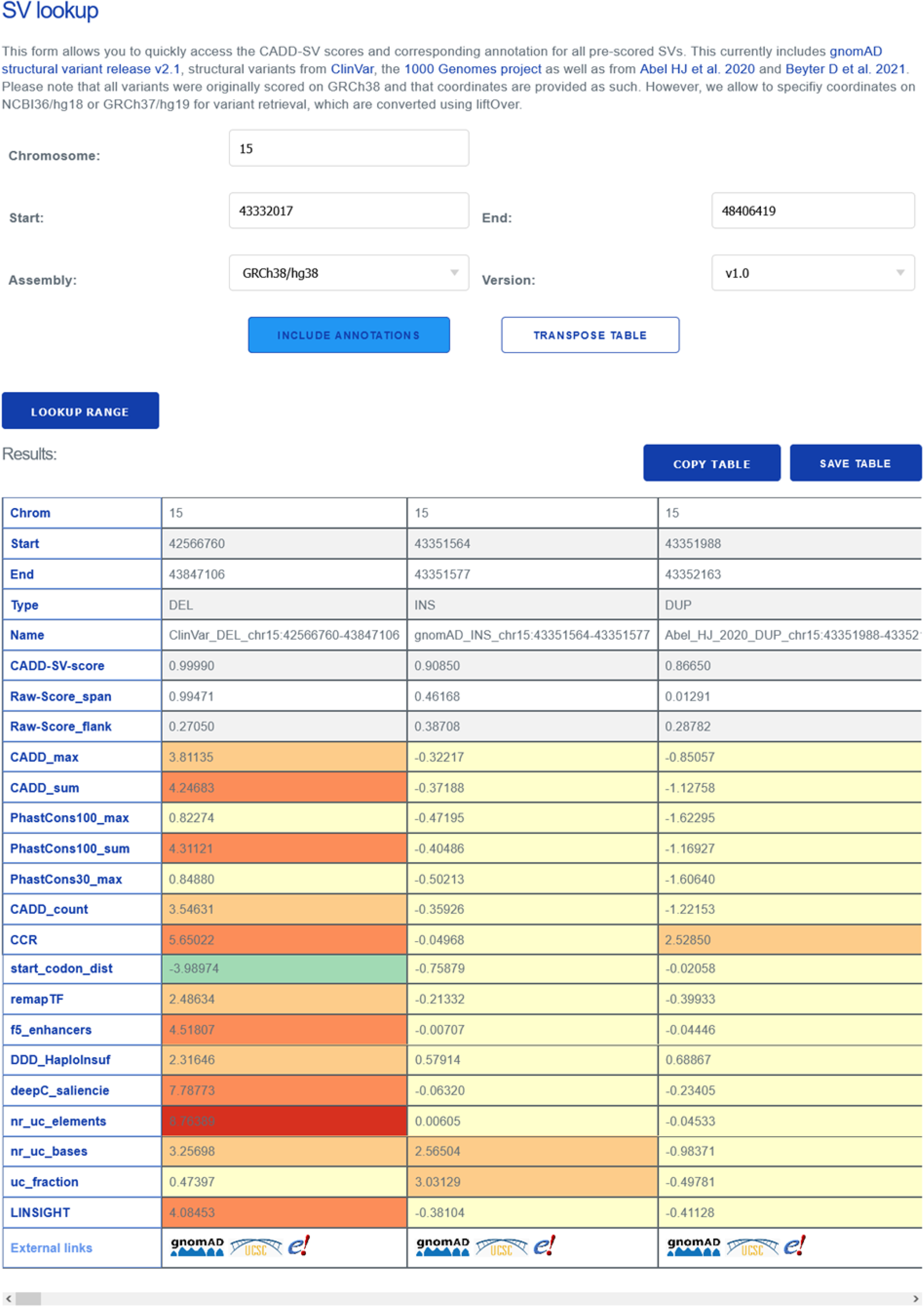
The CADD-SV webserver can score custom SV sets, but it can also be used for direct lookup of pre-scored deletions, duplications and insertions from gnomAD, ClinVar, as well as call-sets from Abel et al. [52] and Beyter et al. [53]. For a given SV, the website provides scores as well as annotations (Z-score) normalized to the value range in the healthy gnomAD cohort. This enables users to infer outliers directly not just by the CADD-SV score but also by color-highlighted annotations. Further, the website provides direct links for each SV to further external resources in gnomAD, Ensembl or the UCSC genome browser.

#### Pathogenic germline variants

We collected pathogenic SVs from ClinVar (n=3262 deletions, 82 duplications and 78 insertions). To look at how CADD-SV prioritizes pathogenic variants among all SVs identified in single individuals (including rare and singleton events), we added each one clinically characterized SV from ClinVar into sets of structural variants found in presumed healthy individuals from the 1000 Genomes project [32]). We assessed CADD-SVs performance by looking at the pathogenic variants rank among all observed SVs. We found that in 50.5% of cases the ClinVar deletion is within the top fifth percentile of all ranks (Fig. 3B). Clinically labelled insertions and duplications were also enriched among the top candidates. In 86% of individuals for insertions and 49% of individuals for duplications do these events fall within the top fifth percentiles.

Further, we contrasted the complete sets of pathogenic SVs from ClinVar with a matched number of common SVs from healthy individuals in gnomAD (AF >= 0.05, Fig. 3D-F). CADD-SV correctly identifies a vast majority of the known pathogenic SVs with an Area Under the ROC Curve (AUROC) of 0.942 for deletions (Fig 3D). CADD-SV performs comparable to the existing SVScore ([20], AUROC of 0.915) and AnnotSV ([22], AUROC of 0.949) methods and outperforms TAD-Fusion score ([21], AUROC of 0.692) in this task.

#### Depletion of deleterious SVs in healthy populations

We assessed the distribution of CADD-SV scores in SVs from healthy individuals of the gnomAD SV call-set. Allele frequency values are significantly decreased in the pathogenic tail of the CADD-SV score distribution compared to the benign tail (top/bottom fifth percentile CADD-SV scores, two-sided Wilcoxon rank sum test, p-value < 10^−16^). We reason that CADD-SV is able to prioritize deleterious variants in healthy individuals as these variants would be under negative selection and removed from the gene pool. Accordingly, the proportion of singleton deletions amongst the top fifth percentile CADD-SV scores (pathogenic tail) is 1.3 times higher than the average of the full SV set (Fig. 3C). This observation is striking for deletions but less pronounced in the insertion and duplication SV sets (Suppl. Figure 9). We note that in the top fifth percentile, 35% of deletions are coding variants classified as “Loss of Function” by gnomAD compared to 0.3 % of variants scored in the remainder of the CADD-SV score distribution.

Further, the average deletion length is six times longer for the top fifth percentile compared to the rest of the distribution, suggesting that longer deletions are more likely to be functional as they affect more sequence. However, short (less than 100bp) and high scoring (top fifth percentile) deletions are 1.1 times more likely to be singletons compared to short deletions, suggesting that CADD-SV prioritizes SVs beyond length. In addition, we detect high frequency deleterious variants in the pathogenic tail, speculating that these variants could be phenotypically functional variants and potentially beneficial for carriers.

#### Identifying somatic cancer variants

We assessed the performance of CADD-SV on somatic variants and the power to identify deleterious cancerogenous variants (n=52,677 deletions, 42,972 duplications and 18 insertions) using SV variants from cancer patients in the International Cancer Genome Consortium [33] as well as insertions reported in Qian et al. [34]. We find an enrichment of SVs detected in cancer patients in the pathogenic tail of the distribution compared to SVs from a healthy cohort (two sided Wilcoxon rank sum test, p-value <10^−16^). CADD-SV enriches the cancer-derived SVs from common gnomAD-SVs in a ROC Curve analysis (AUROC 0.848 / 0.933 / 0.975 for deletions / duplications / insertions, Fig. 3D-F), outperforming available tools on this task and supporting the claim that CADD-SV prioritizes functional somatic SVs.

#### Identifying expression altering non-coding variants

To test the ability to prioritize functional variants beyond coding regions, we use a set of non-coding SVs known to alter the expression of genes. Here, we look at 387 deletions and 300 duplications that were shown to affect expression levels of nearby genes and are therefore considered expression Quantitative Trait Loci (eQTL) by the GTEx consortium [1]. We compare them against common variants (AF >= 0.05) from gnomAD in a ROC Curve analysis (Fig. 3D-F). Even though less pronounced compared to ClinVar or the cancer-derived SVs, CADD-SV is able to differentiate the two classes of SVs (AUROC 0.598 for deletions and 0.635 for duplications, respectively) outperforming existing methods SVScore (AUROC 0.467 / 0.534), AnnotSV (AUROC 0.459 / 0.402) and TAD-Fusion score (AUROC 0.425 for deletions).

### Interpreting Structural Variants

To make scores easier to interpret, we rank CADD-SV raw scores among 20,000 SVs from healthy individuals in gnomAD. Final CADD-SV scores range from 0 (potentially benign) to 1 (potentially pathogenic), indicating the position of the novel variant within the gnomAD score distribution. For example, a value of 0.9 represents that 90% of variants reported from healthy individuals are scoring lower than the variant under consideration. In addition, all feature annotations are used and reported after Z-score transformation (mean 0, standard deviation of 1) according to the features value distribution in gnomAD. This allows users to inspect the individual features for extreme values easily. For instance, a conservation feature value of four represents an outlier value of four standard deviations away from the gnomAD mean of that annotation. Such noticeable values are highlighted by color-coding on the CADD-SV website (Figure 4) for the pre-scored variant sets. Generally, CADD-SV scores with or without annotation information are available from our command line tool as well as on the webserver for direct variant interpretation. Our online services include region lookups of existing SV datasets, coordinate transfers between human genome builds, the download of pre-scored datasets and annotations, a simple API for pre-scored variants as well as the online scoring of novel SV datasets. Coordinate ranges and variants of other genome builds (i.e. GRCh37/hg19 and NCBI36/hg18) can be used on the webserver and are automatically lifted to GRCh38 coordinates (providing the original coordinates in the variant’s name field).

## Discussion

We present CADD-SV as an unbiased and powerful tool for the annotation and prioritization of deleterious structural variants. CADD-SV is built from machine learning models that are trained using evolutionary-derived and putative benign variants that underwent millions of years of purifying selection. These variants are contrasted with a background set of the same size and length, encountering deleterious events by chance. We show that our approach is able to model and score deletions, insertions as well as duplication and we validate the CADD-SV models using clinically annotated, non-coding or population germline SVs as well as somatic SVs reported in cancer patients.

Structural variant calling is prone to biases towards certain types of SVs, as the signal to for example detect deletions is vastly different compared to signals of duplication or even inversions [19]. Further, the exact annotation of SV breakpoints is often limited, e.g. due to their frequent positioning in repetitive sequence [46]. Apart from these universal limitations, changes in the application of arrays and sequencing technologies over the last decades have affected available SV sets. However, in previous work it seems underappreciated how much the historic and functional ascertainment imprinted on potential training and validation sets for machine learning. Specifically, the ClinVar-annotated SVs are comparably large and clustered around well-studied genes. Using an alternative source for the training data, the CADD-SV approach is not confounded and performance can be evaluated broadly, as allele-frequency features or any ClinVar annotations are not included in the features or otherwise considered when building the training sets. The number of labelled SVs to validate the performance of CADD-SV is still limited though. Assessing the performance on duplications and insertions is particularly hard as the number of known pathogenic events is small and strongly biased towards coding sequence. We anticipate that future datasets will provide a better opportunity to test and interpret models for duplications and insertions.

Estimating functional effects of SVs is highly complex due their size (involving different molecular targets) but also due to different mechanistic types of SVs (e.g. deletion, insertion, duplication or inversion of sequence). Thus, deleteriousness effects cannot just result from the sequence alteration, but also from interactions with the sequence context. For example, sequences shielding gene regulation (e.g. TAD boundaries) can be deleted between coding sequences or non-functional sequence can be inserted, interfering with an existing regulatory unit. Therefore, we model each SV type (deletions, insertions and duplications) separately, and we use the sequence span as well as the flanking sequence regions to capture putative pathogenic effects comprehensively. Further, we integrate distance features and a large set of annotations covering both coding and non-coding effects. This allows CADD-SV high predictive performance on known disease variants from ClinVar, which often cover coding sequence and stand-out by their gene model annotations and genes scores such as pLI [47] or DDD Haploinsufficiency [48]. Further, putative pathogenic non-coding variants can be prioritized using sequence conservation [42], enhancer annotations [49,50] and links [51], assay readouts such as RNase-seq or ChIP-seq, as well as information about 3D interactions from the Hi-C directionality index [43,44] or computational predictions such as deepC [45]. These kinds of mechanisms were previously shown to be causal for human disease phenotypes [6].

Inversions and translocations are particularly hard to assess as they are copy number neutral and their impact is often mediated by proximity of certain functional elements to one another or functional entities such as TADs being broken or reshuffled rather than deleting or inserting functional sequence directly. To our knowledge, there is no training dataset sufficient in size and curation to capture the complexity of these events. As no single model captures the mechanistic diversity of the currently considered types of structural variants, CADD-SV reports normalized model scores and features, normalized to the same type of SVs identified from healthy individuals. Feature normalization enables users to inspect annotation outliers directly (reported as standard deviation away from the mean), visually highlighting certain annotations and hinting at potential pathogenic mechanisms beyond the CADD-SV score.

In contrast to other tools, length is not a feature of CADD-SV. However, we assume that SV length would be a good indicator of SV impact, as long SVs are more likely to affect coding regions or generally functional annotations. SV length itself might be a confounder too, as long benign SVs might be misinterpreted solely for their length and not for their actual genomic signatures. As the contrasting datasets in the CADD-SV framework are matched in SV length, length as a feature does not contribute to the model. However, some genomic feature transformation such as the sum of all intersected annotation values or the number of bases above a certain threshold, correlate inevitably with length but are bound to functional annotations being present across the span. AnnotSV [22] is powerful and efficient in annotating novel SVs with a wide set of annotations. However, validation of AnnotSV on ClinVar is biased as AnnotSV uses overlap of novel SVs with labelled SVs from ClinVar as a feature. Further, it categorizes SVs in five bins from benign to pathogenic instead of a continuous score. Across multiple data sets, we highlight the increased predictive power of CADD-SV compared to AnnotSV, SVscore [20]and TAD-Fusion [21]. A comparison of SVFX [23]was not possible, as the package is not easily deployed, explicitly normalizes features on each training data set and its released ClinVar variant scores are based on a model trained on the same variant set (a biased comparison in which it outperforms our scores; data not shown).

The feature integration implemented by CADD-SV can easily be extended using additional annotations. Currently, we use features derived from experiments conducted in specific cell-types (e.g. GM12878, H1, A549, CAKI2). More comprehensive or additional cell-types can be included in updated versions. Further, CADD-SV does not make us of the inserted sequence itself. Therefore, future versions of CADD-SV could make use of sequence-based prediction models in addition to reference annotations, e.g. to predict open reading frames, repeat content, presence of transcription factor binding sites or the general likelihoods of the novel inserted sequence being of open or closed chromatin. This might be powerful in assessing inserted sequence function beyond the surrounding genomic context of the insertion event. In addition, specific mechanistic events such as gene-fusion predictions are not part of our features and CADD-SV is therefore only able to estimate the effect of those events based on other already considered feature values like the distance to genes.

CADD-SV integrates rich sets of annotations in predictive models of SV effects for deletions, insertions and duplications. It is designed for genome build GRCh38 but can be applied to other genome builds due to an integrated liftover step. The output consists of a comprehensive score of deleteriousness with higher values corresponding to larger effect variants. We normalize CADD-SV scores on germline SVs from healthy individuals and report the percentile for a novel SV within this score distribution. CADD-SV provides all integrated feature values as an output table for users to screen the predicted effects for gene overlap or various other functional effects.

## Data Availability

CADD-SV pre-scored variant sets as well as a webtool for the interpretation of novel deletions, insertions and duplications is available (https://cadd-sv.bihealth.org/). The CADD-SV framework can be cloned and used from GitHub (https://github.com/kircherlab/CADD-SV/). All external data sets used are available under the locations specified in the Methods. Further information on the analyses is available on request.

## Acknowledgments

We thank current and previous members of the Kircher lab for helpful discussions and suggestions. Specifically, we would like to acknowledge Kunaphas Kongkitimanon for his contributions to the website as well as Lusine Nazaretyan for feedback on the GitHub manual. Computation has been performed on the HPC for Research cluster of the Berlin Institute of Health.

## References

1. Chiang C, Scott AJ, Davis JR et al. The impact of structural variation on human gene expression. Nat Genet 2017;49:692–9.

2. Collins RL, Brand H, Karczewski KJ et al. A structural variation reference for medical and population genetics. Nature 2020;581:444–51.

3. Lupiáñez DG, Kraft K, Heinrich V et al. Disruptions of Topological Chromatin Domains Cause Pathogenic Rewiring of Gene-Enhancer Interactions. Cell 2015;161:1012–25.

4. Sudmant PH, Rausch T, Gardner EJ et al. An integrated map of structural variation in 2,504 human genomes. Nature 2015;526:75–81.

5. Rodriguez-Revenga L, Mila M, Rosenberg C et al. Structural variation in the human genome: the impact of copy number variants on clinical diagnosis. Genetics in Medicine 2007;9:600–6.

6. Spielmann M, Lupiáñez DG, Mundlos S. Structural variation in the 3D genome. Nature Reviews Genetics 2018:1.

7. Gloss BS, Dinger ME. Realizing the significance of noncoding functionality in clinical genomics. Experimental & Molecular Medicine 2018;50:1–8.

8. Lupiáñez DG, Kraft K, Heinrich V et al. Disruptions of topological chromatin domains cause pathogenic rewiring of gene-enhancer interactions. Cell 2015;161:1012–25.

9. Lupiáñez DG, Spielmann M, Mundlos S. Breaking TADs: How Alterations of Chromatin Domains Result in Disease. Trends Genet 2016;32:225–37.

10. Lieberman-Aiden E, van Berkum NL, Williams L et al. Comprehensive mapping of long-range interactions reveals folding principles of the human genome. Science 2009;326:289–93.

11. Gasperini M, Tome JM, Shendure J. Towards a comprehensive catalogue of validated and target-linked human enhancers. Nature Reviews Genetics 2020:1–19.

12. The ENCODE Project Consortium. An integrated encyclopedia of DNA elements in the human genome. Nature 2012;489:57–74.

13. Inoue F, Ahituv N. Decoding enhancers using massively parallel reporter assays. Genomics 2015;106:159–64.

14. Kircher M, Xiong C, Martin B et al. Saturation mutagenesis of twenty disease-associated regulatory elements at single base-pair resolution. Nat Commun 2019;10:1–15.

15. Nguyen TA, Jones RD, Snavely AR et al. High-throughput functional comparison of promoter and enhancer activities. Genome Res 2016;26:1023–33.

16. Santiago-Algarra D, Dao LTM, Pradel L et al. Recent advances in high-throughput approaches to dissect enhancer function. F1000Res 2017;6, DOI: 10.12688/f1000research.11581.1.

17. Haller F, Bieg M, Will R et al. Enhancer hijacking activates oncogenic transcription factor NR4A3 in acinic cell carcinomas of the salivary glands. Nature Communications 2019;10:368.

18. Helmsauer K, Valieva ME, Ali S et al. Enhancer hijacking determines extrachromosomal circular MYCN amplicon architecture in neuroblastoma. Nature Communications 2020;11:5823.

19. Cameron DL, Di Stefano L, Papenfuss AT. Comprehensive evaluation and characterisation of short read general-purpose structural variant calling software. Nature Communications 2019;10:3240.

20. Ganel L, Abel HJ, Hall IM. SVScore: an impact prediction tool for structural variation. Bioinformatics 2017;33:1083–5.

21. Huynh L, Hormozdiari F. TAD fusion score: discovery and ranking the contribution of deletions to genome structure. Genome Biology 2019;20:60.

22. Geoffroy V, Herenger Y, Kress A et al. AnnotSV: an integrated tool for structural variations annotation. Bioinformatics 2018;34:3572–4.

23. Kumar S, Harmanci A, Vytheeswaran J et al. SVFX: a machine-learning framework to quantify the pathogenicity of structural variants. bioRxiv 2019:739474.

24. Landrum MJ, Lee JM, Benson M et al. ClinVar: improving access to variant interpretations and supporting evidence. Nucleic Acids Res 2018;46:D1062–7.

25. Kronenberg ZN, Fiddes IT, Gordon D et al. High-resolution comparative analysis of great ape genomes. Science 2018;360, DOI: 10.1126/science.aar6343.

26. Quinlan AR, Hall IM. BEDTools: a flexible suite of utilities for comparing genomic features. Bioinformatics 2010;26:841–2.

27. Köster J, Rahmann S. Snakemake—a scalable bioinformatics workflow engine. Bioinformatics 2012;28:2520–2.

28. Liaw A, Wiener M. Classification and Regression by randomForest. R News 2002;2:18–22.

29. Grau J, Grosse I, Keilwagen J. PRROC: computing and visualizing precision-recall and receiver operating characteristic curves in R. Bioinformatics 2015;31:2595–7.

30. Hancks DC, Kazazian HH. Active human retrotransposons: variation and disease. Current Opinion in Genetics & Development 2012;22:191–203.

31. Gardner EJ, Prigmore E, Gallone G et al. Contribution of retrotransposition to developmental disorders. Nature Communications 2019;10:4630.

32. Auton A, Abecasis GR, Altshuler DM et al. A global reference for human genetic variation. Nature 2015;526:68–74.

33. Campbell PJ, Getz G, Korbel JO et al. Pan-cancer analysis of whole genomes. Nature 2020;578:82–93.

34. Qian Y, Mancini-DiNardo D, Judkins T et al. Identification of pathogenic retrotransposon insertions in cancer predisposition genes. Cancer Genetics 2017;216–217:159–69.

35. Kuhn RM, Haussler D, Kent WJ. The UCSC genome browser and associated tools. Brief Bioinform 2013;14:144–61.

36. Grüning B, Dale R, Sjödin A et al. Bioconda: sustainable and comprehensive software distribution for the life sciences. Nat Methods 2018;15:475–6.

37. Huang Y-F, Gulko B, Siepel A. Fast, scalable prediction of deleterious noncoding variants from functional and population genomic data. Nat Genet 2017;49:618–24.

38. Rentzsch P, Witten D, Cooper GM et al. CADD: predicting the deleteriousness of variants throughout the human genome. Nucleic Acids Res 2019;47:D886–94.

39. Collins RL, Brand H, Karczewski KJ et al. An open resource of structural variation for medical and population genetics. bioRxiv 2019:578674.

40. Kircher M, Witten DM, Jain P et al. A general framework for estimating the relative pathogenicity of human genetic variants. Nature Genetics 2014;46:310–5.

41. Davydov EV, Goode DL, Sirota M et al. Identifying a High Fraction of the Human Genome to be under Selective Constraint Using GERP++. Wasserman WW (ed.). PLoS Computational Biology 2010;6:e1001025.

42. Siepel A, Bejerano G, Pedersen JS et al. Evolutionarily conserved elements in vertebrate, insect, worm, and yeast genomes. Genome Res 2005;15:1034–50.

43. Calandrelli R, Wu Q, Guan J et al. GITAR: An Open Source Tool for Analysis and Visualization of Hi-C Data. Genomics, Proteomics & Bioinformatics 2018;16:365–72.

44. Schmitt AD, Hu M, Jung I et al. A Compendium of Chromatin Contact Maps Reveals Spatially Active Regions in the Human Genome. Cell Reports 2016;17:2042–59.

45. Schwessinger R, Gosden M, Downes D et al. DeepC: predicting 3D genome folding using megabase-scale transfer learning. Nature Methods 2020:1–7.

46. Kosugi S, Momozawa Y, Liu X et al. Comprehensive evaluation of structural variation detection algorithms for whole genome sequencing. Genome Biology 2019;20:117.

47. Lek M, Karczewski KJ, Minikel EV et al. Analysis of protein-coding genetic variation in 60,706 humans. Nature 2016;536:285–91.

48. Firth HV, Wright CF. The Deciphering Developmental Disorders (DDD) study. Developmental Medicine & Child Neurology 2011;53:702–3.

49. Abugessaisa I, Noguchi S, Hasegawa A et al. FANTOM5 CAGE profiles of human and mouse reprocessed for GRCh38 and GRCm38 genome assemblies. Sci Data 2017;4:170107.

50. Chèneby J, Gheorghe M, Artufel M et al. ReMap 2018: an updated atlas of regulatory regions from an integrative analysis of DNA-binding ChIP-seq experiments. Nucleic Acids Research 2017, DOI: 10.1093/nar/gkx1092.

51. Hait TA, Amar D, Shamir R et al. FOCS: a novel method for analyzing enhancer and gene activity patterns infers an extensive enhancer-promoter map. Genome Biol 2018;19:56.

52. Abel HJ, Larson DE, Chiang C et al. Mapping and characterization of structural variation in 17,795 deeply sequenced human genomes. bioRxiv 2018:508515.

53. Beyter D, Ingimundardottir H, Oddsson A et al. Long-read sequencing of 3,622 Icelanders provides insight into the role of structural variants in human diseases and other traits. Nature Genetics 2021:1–8.

