## Supplementary Figures for "CADD-SV – a framework to score the effects of structural variants in health and disease"

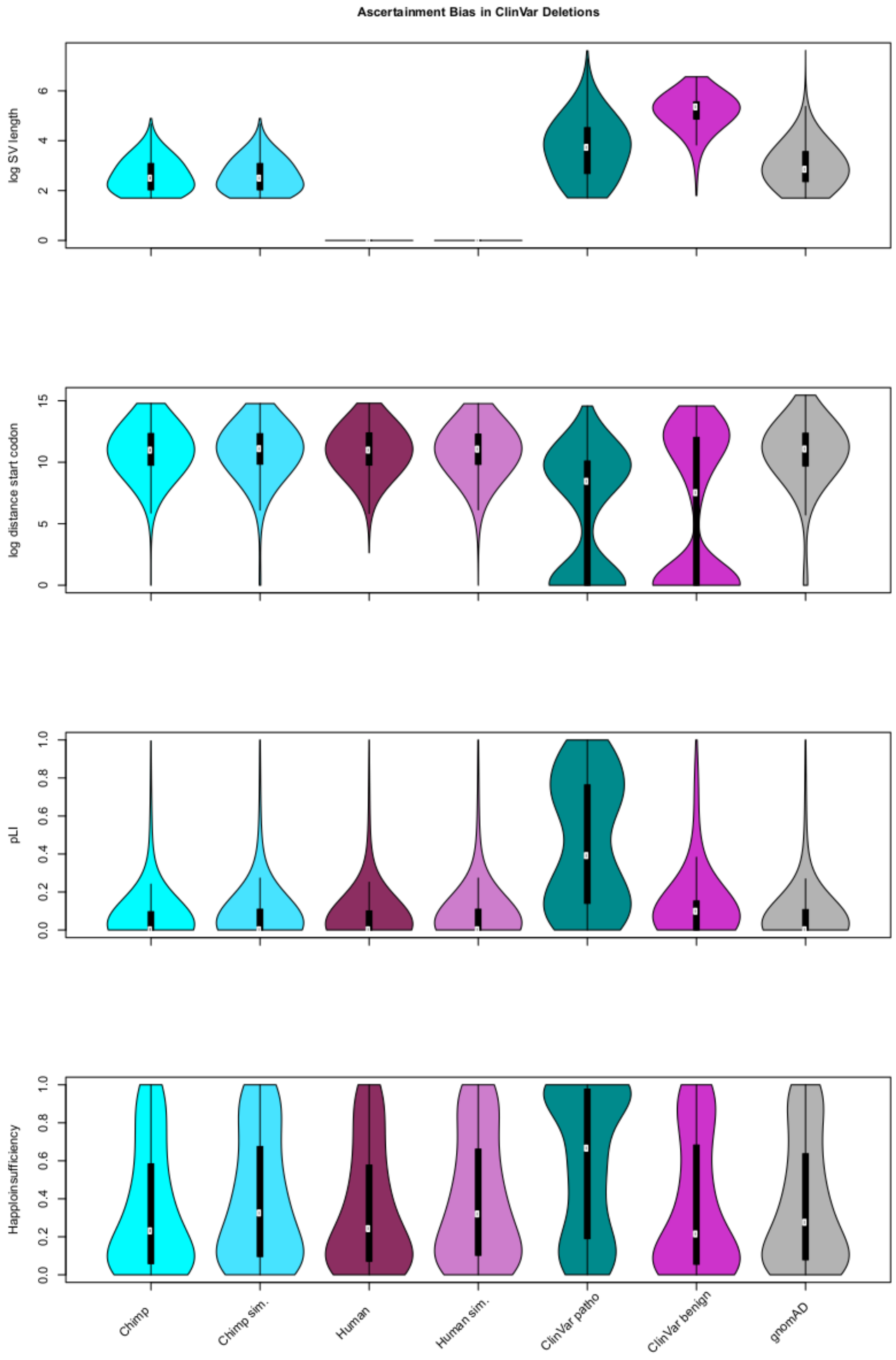


***Supplemental Figure 1:*** *Ascertainment bias in labeled deletion datasets. To make accurate predictions using machine learning it is crucial to have an unbiased dataset to train on. ClinVar pathogenic or benign labelled deletions are hand curated and individually verified but are biased towards very large deletions and are clustering around well-studied genes (as shown in the excess of high pLI and Haploinsufficiency scores). Our evolutionary derived dataset however does not suffer from these kinds of ascertainment bias and is similar to the occurrence of deletions in a healthy population cohort (gnomAD-SV v2.0).*

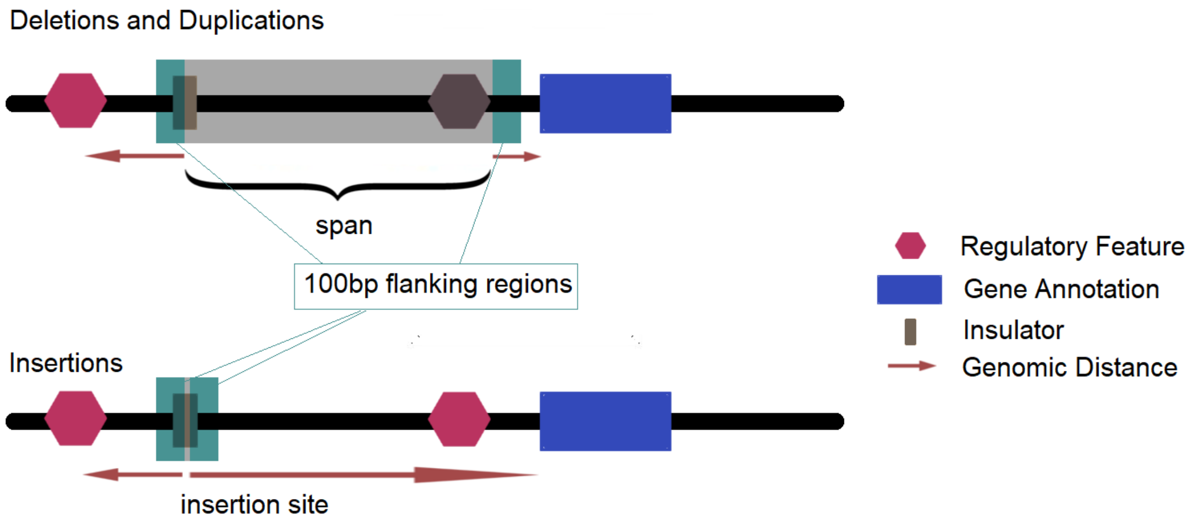


***Supplemental Figure 2:*** *Deletions and duplications stretch genomic sequence that can be annotated with comprehensive features. Novel insertions however are only annotated as the site of integration. In addition, all types of SVs are annotated over 100bp down- and upstream sequence of the variant. Further distance features to the closest gene/regulatory feature are collected for span and flank models. We train the span of novel deletions with the chimp DEL set and train the sequence 100bp up- and downstream of the breakpoints using the human DEL set. We use the chimp INS set for the insertion site and the human INS set for the up- and downstream sequence. Duplications are scored using the full sequence span of the duplicated locus, hence using the chimp DEL model for the span and human DEL model for the up- and downstream sequence. The final score is calculated from the maximum (more deleterious) value of both models applied after being ranked compared to the score distribution of the same type of SVs reported in healthy individuals.*

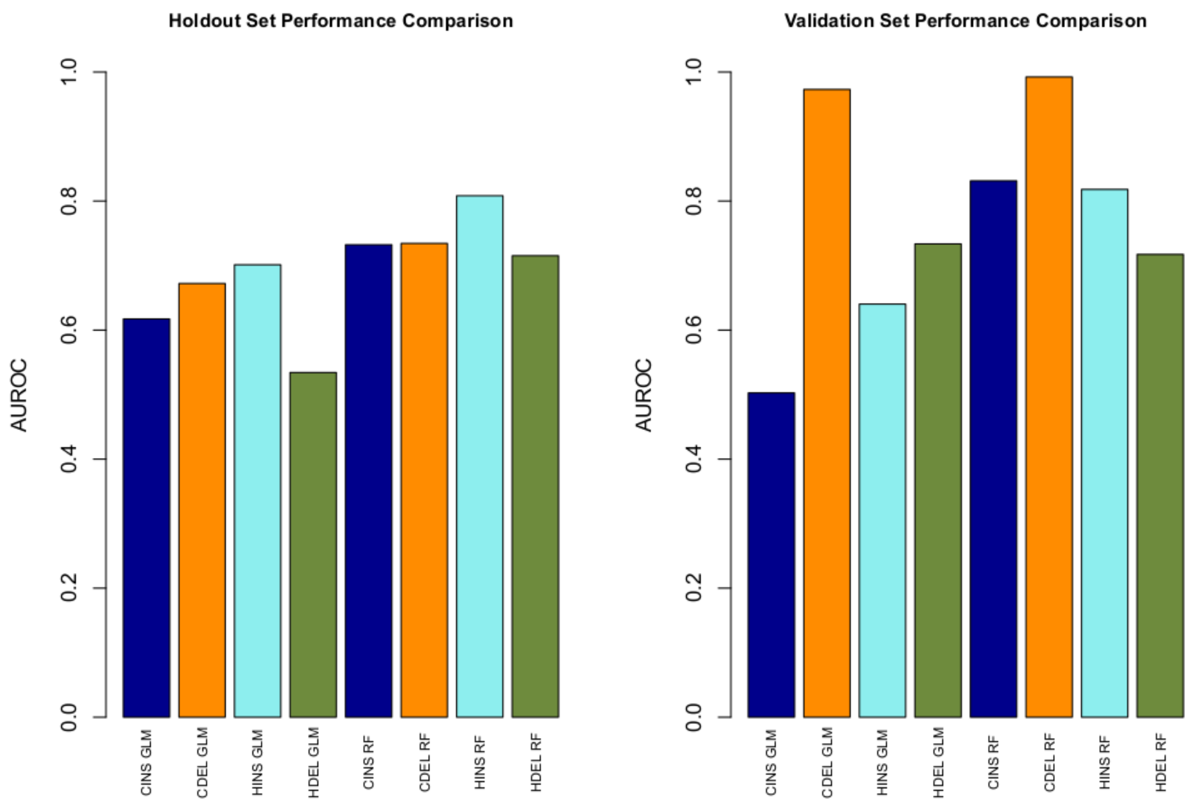


***Supplemental Figure 3:*** *Model comparison of Random Forrest classifiers and generalized linear models trained using the R GLM package. We validated the performance of both classifiers using 10% randomly sampled holdout data (left) as well as one of the validation sets (labelled pathogenic SVs from ClinVar vs. SVs in gnomAD, right).*

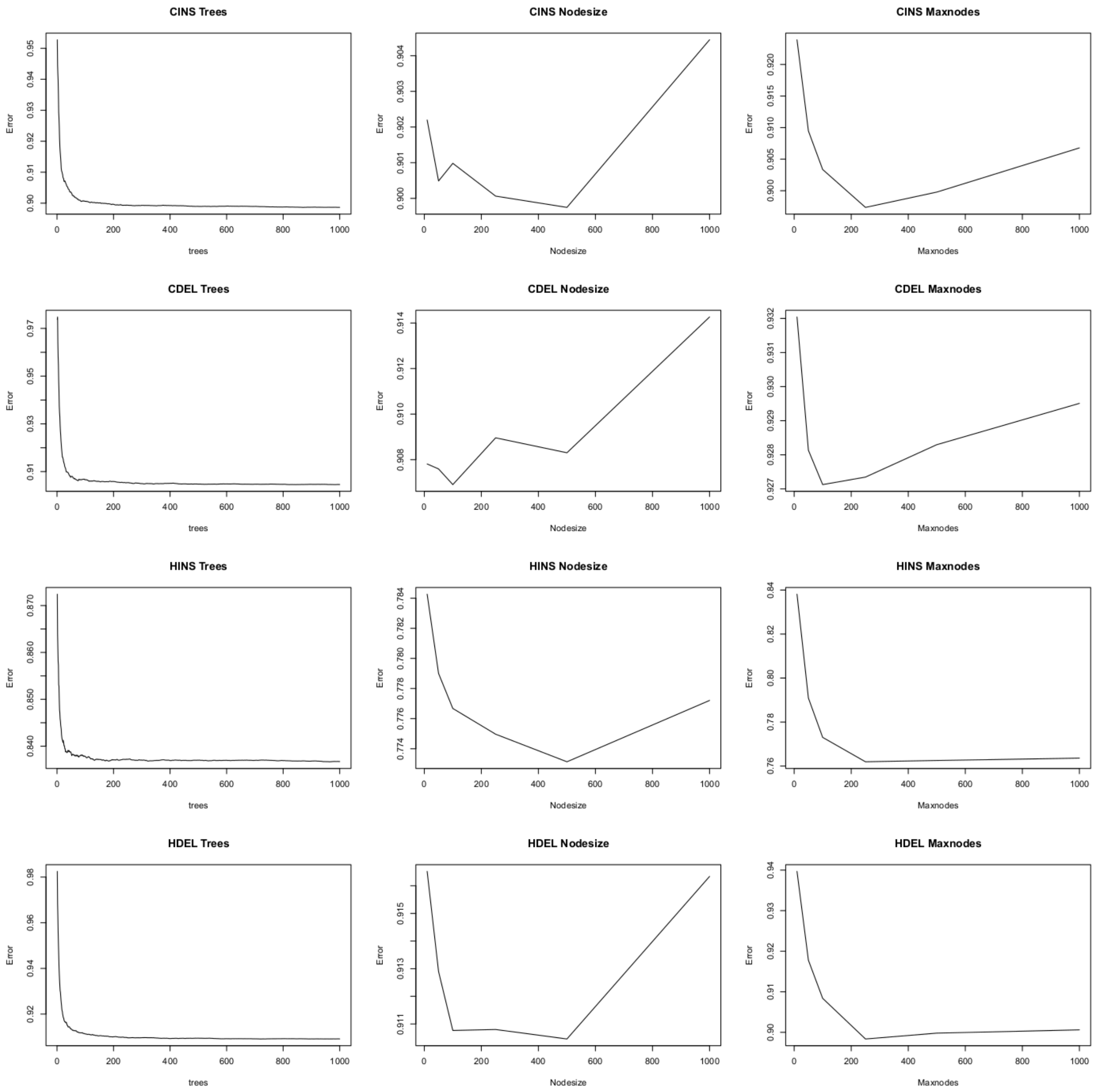


***Supplemental Figure 4:*** *Hyperparameter search for all Random Forest models. We explored the mean of squared errors for the parameters nodesize and maxnodes for both values of n= {10, 50, 100, 250, 500, 1000} and chose parameters minimizing error and overfitting. The number of trees (ntree = {25, 50, 75, 100, 200, 500, 1000}) for each model was chosen based on observing no further improvement of error by increasing the number of trees. CINS: Chimp Insertions, CDEL: Chimp Deletions, HINS: Human Insertions, HDEL: Human Deletions*

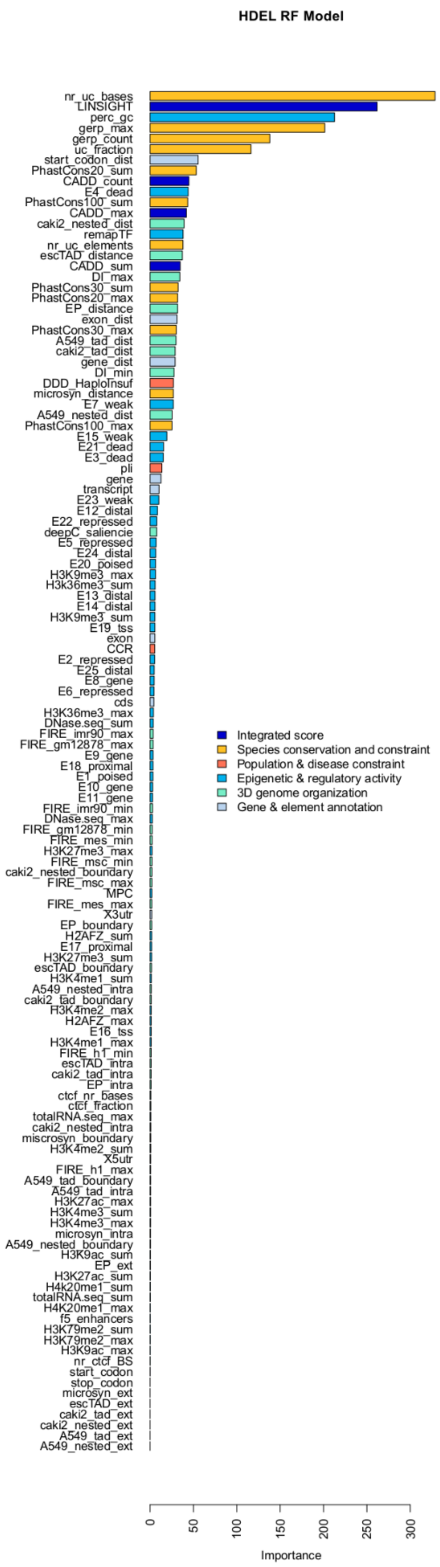


***Supplemental Figure 5:*** *Feature contribution of human deletion (human DEL) flank model.*

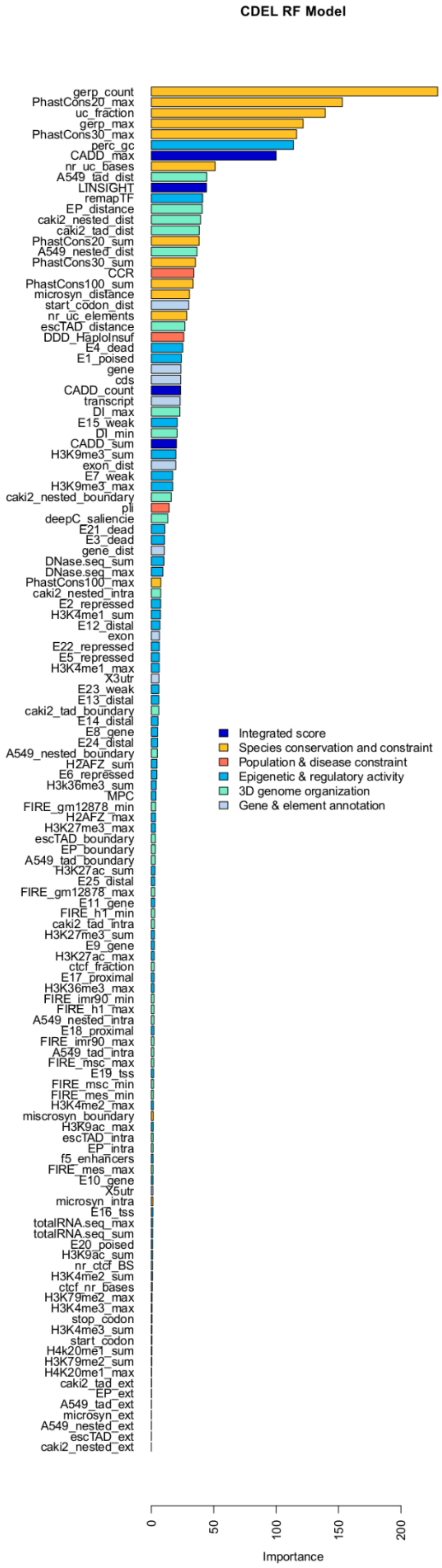


***Supplemental Figure 6:*** *Feature contribution of chimpanzee deletion (chimp DEL) span model.*

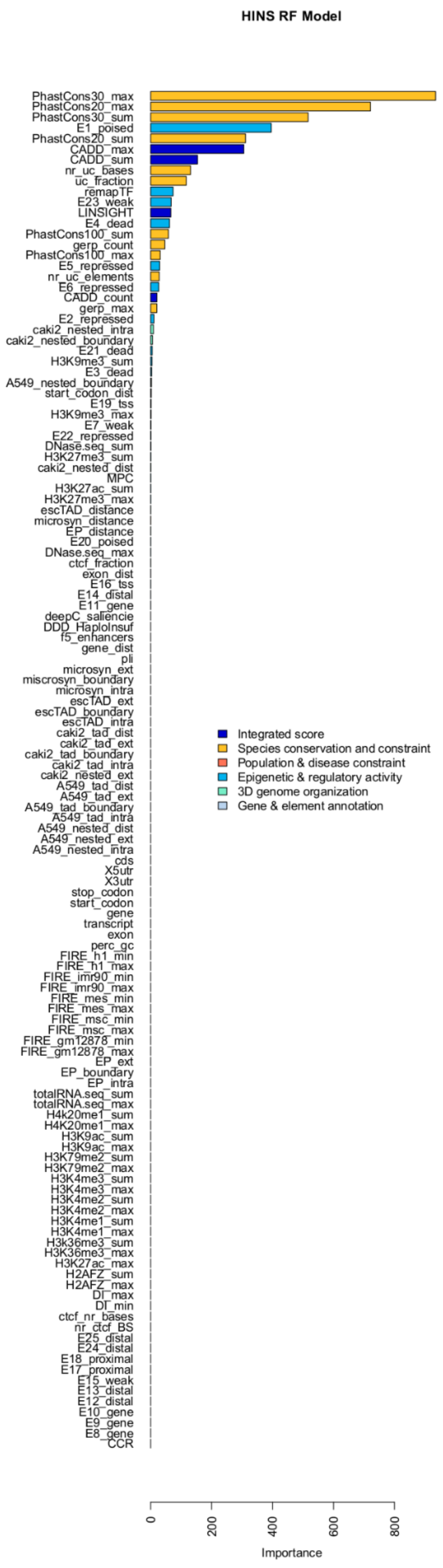


***Supplemental Figure 7:*** *Feature contribution of the human insertion (human INS) flank model.*

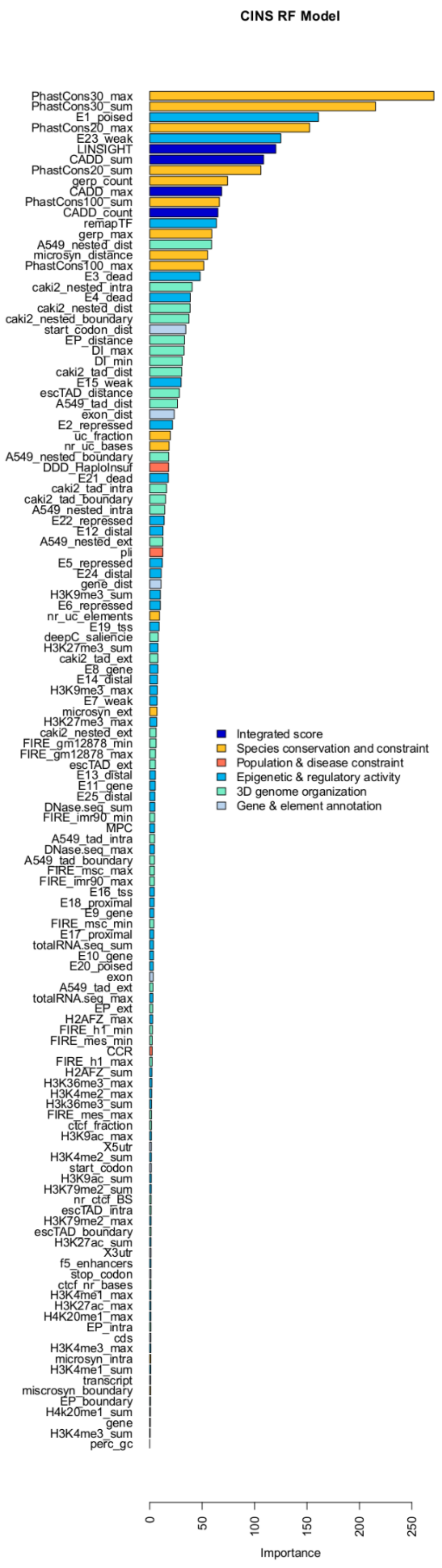


***Supplemental Figure 8:*** *Feature contributions of chimpanzee insertion (chimp INS) span/site model.*

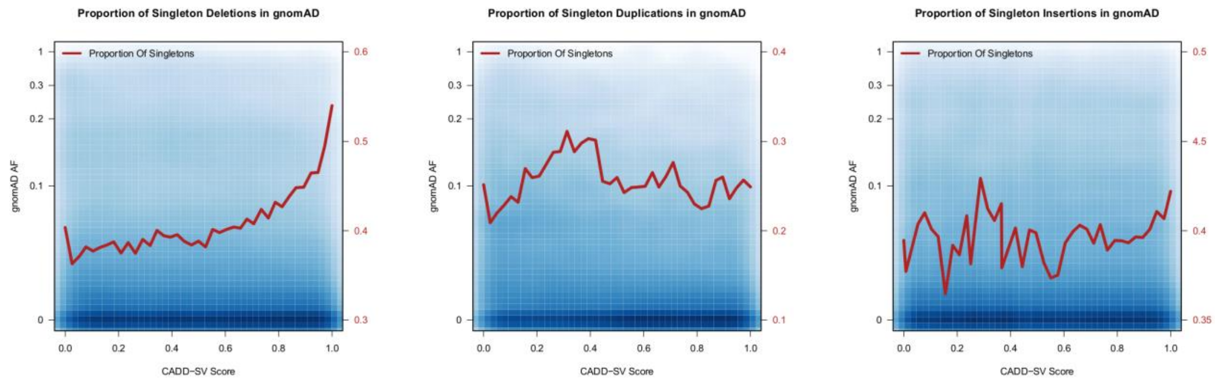


***Supplemental Figure 9:*** *Proportion of singleton insertions and duplications in the gnomAD-SV data set of putative healthy individuals. The pathogenic CADD-SV score tail (1 being most pathogenic) is enriched in singletons, suggesting purifying selection against SVs with high CADD-SV scores. However, this effect is less pronounced in insertions and duplications compared to the deletions.*

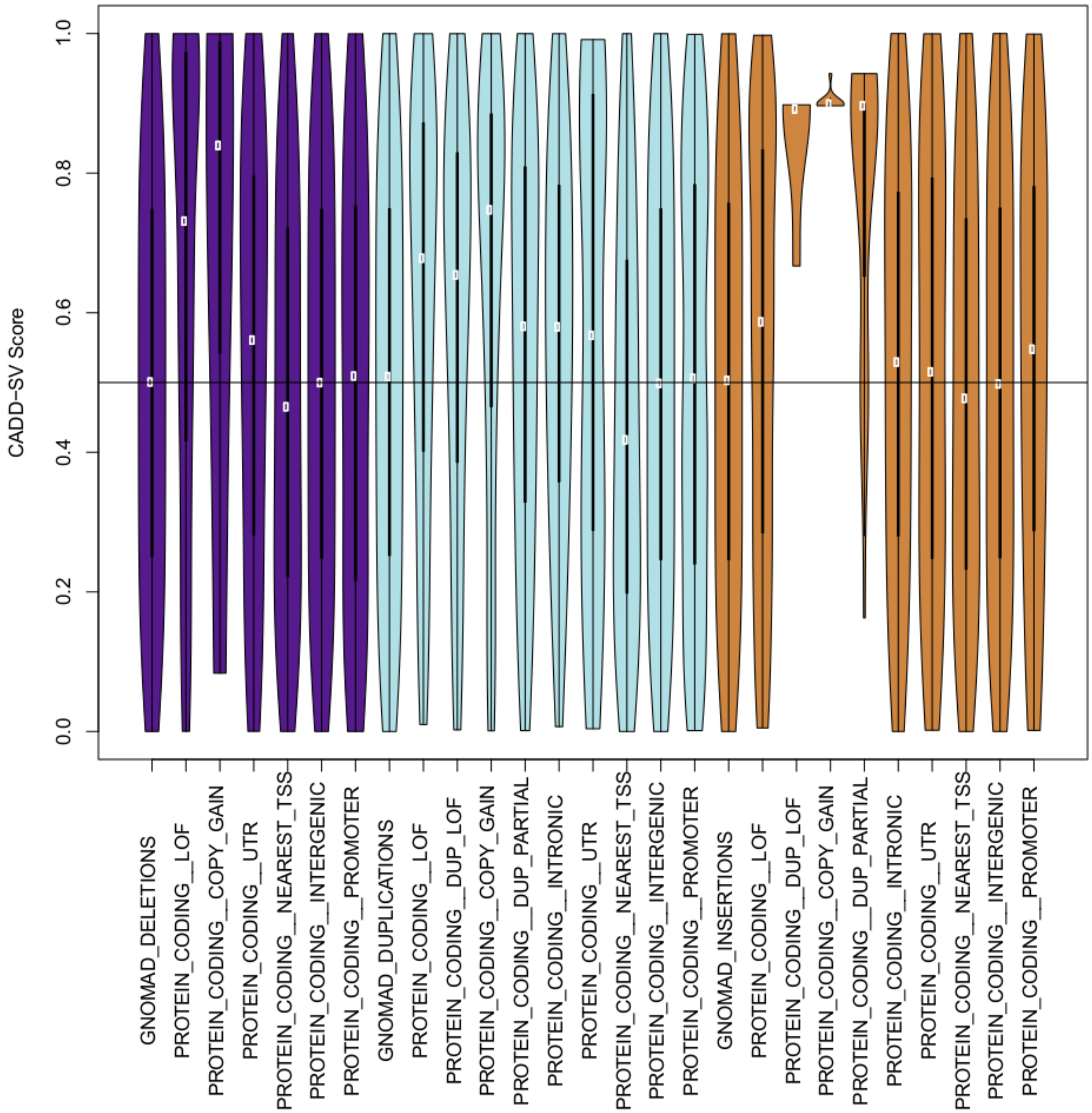


***Supplemental Figure 10:*** *CADD-SV Score distribution for gnomAD deletions (purple), duplications (light blue) and insertions (orange) for various categories annotated in the gnomAD-SV release. Increased CADD-SV score suggest increased effect sizes.*
